# Game theory of vaccination and depopulation for managing avian influenza on poultry farms

**DOI:** 10.1101/348813

**Authors:** Alexis Delabouglise, Maciej F Boni

**Author notes:** **Correspondence** Alexis Delabouglise Millennium Sciences Complex Department of Biology Pennsylvania State University University Park, PA, 16802.

## Abstract

Highly pathogenic avian influenza is endemic in domestic poultry populations in East and South Asia and is a major threat to human health, animal health, and the poultry production industry. The behavioral response of farmers to the disease and its epidemiological effects are still poorly understood. We considered a symmetric game in a region with widespread smallholder poultry production, where the players are broiler poultry farmers and between-farm disease transmission is both environmental (local) and mediated by the trade of infected birds. Three types of farmer behaviors were modelled: vaccination, depopulation, and cessation of poultry farming. We found that the transmission level of avian influenza through trade networks had strong qualitative effects on the system’s epidemiological-economic equilibria. In the case of low trade-based transmission, when the monetary cost of infection is sufficiently high, depopulation behavior persists and maintains a disease-free equilibrium. In the case of high trade-based transmission, depopulation behavior has perverse epidemiological effects - as it accelerates the spread of disease via poultry traders - but has a high enough payoff to farmers that it persists at the system’s game theoretic equilibrium. In this situation, state interventions should focus on making effective vaccination technologies available at a low price rather than penalizing infected farms. Our results emphasize the need in endemic countries to further investigate the commercial circuits through which birds from infected farms are traded.

## Introduction

Behavioral epidemiology - the study of human behavioral responses to infectious disease circulation - has been receiving an increasing amount of research attention over the last two decades (1). The behavioral responses of humans to disease can generate unexpected externalities resulting in both positive and negative feedback in the infection process, justifying the application of game theory to questions in the decentralized control of infectious disease. Examples of theoretical advances in this field have focused on the adoption of voluntary vaccination programs (2), the effects of endogenous vs government-recommended social distancing strategies (3), and the willingness of states to disclose information on disease outbreaks (4). A recent empirical study, based on time series of reports of social contacts, confirmed that voluntary social distancing behavior affected the dynamics of H1N1 influenza epidemics in the US (5).

Although the effects of human behavior on the epidemiology of livestock disease are well-documented, empirical studies on the effects of disease on human behavior are rare (6, 7). In livestock disease epidemiology, it is reasonable to assume that producers primarily aim to maximize their protein production or revenue and minimize their farm expenses; this principle may be used to predict livestock owners’ behavioral response to disease risk. From a social planner’s perspective an additional optimization is centered on the choice of public intervention, balancing the social need for affordable poultry products, the welfare of actors in livestock value chains, and protecting public health. Therefore, any particular surveillance and control policy must be evaluated by comparing its associated gains in public health with its effect on producers’ income and consumers’ access to livestock products.

Highly pathogenic avian influenza (HPAI) is a zoonotic livestock disease motivating substantial state intervention. HPAI outbreaks have occurred regularly in eastern and southern Asia, Egypt, and West Africa since 2003 (8–10). HPAI has also been reported in Europe and North America (9). Some avian influenza virus strains of the H5, H7, and H9 subtypes have the ability to cause severe and fatal disease in humans. Therefore, poultry originating from farms contaminated with HPAI are potentially harmful to farmers, consumers, and other persons handling poultry. While human infections with these subtypes are rare, their case fatality rates are generally higher than 25% (11). In addition, the risk that such viruses acquire a phenotype of human-to-human transmission constitutes a major global health threat and justifies national-level interventions to reduce the exposure of humans to infected poultry (12). Since the emergence and global spread of the H5N1 subtype of HPAI in 2003, interventions have mainly focused on strengthening avian disease surveillance, preventive culling of domestic birds in outbreak areas, and restrictions on poultry trade; in addition, Vietnam and China implemented mandatory poultry vaccination programs (13, 14). A socio-economic field study conducted in Vietnam showed that farmers fear HPAI detection by the surveillance system not so much because of the risk of mandatory culling but because of its adverse effect on the sale price of poultry (15).

In a theoretical model investigating individual responses of poultry farmers to price fluctuations, the main finding for density-dependent transmission was that preventive culling of and compensation for poultry can have perverse effects as it may incentivize an increase in farm size (higher average poultry price incentivizes more production), while investments in surveillance, diagnostics, and targeted penalties were found to lower overall health risk (for both frequency- and density-dependent transmission) (16). These findings are in agreement with general economic models describing moral hazards and adverse selection issues associated with the indemnification of farms affected by diseases (17–19). According to another economic model applied to HPAI, compensation accompanying preventive culling policies should be indexed to the prevention effort invested by farmers (20). These studies focus on the individual responses of farmers and, as such, do not account for population-level externalities (e.g. between-farm infection risk or market price). According to a generic game theory model focusing on prevention practices, enforcing penalties on infected farms seems to be an appropriate way of incentivizing disease control in the private sector, while policies based on indemnification may discourage prevention efforts (19).

The present study aims at modeling the behavioral responses of a heterogeneous group of poultry farmers under different economic and epidemiological conditions. We introduce a symmetric population game for this purpose, with broiler poultry farmers as players, and we link this to a compartmental model of between-flock disease transmission.

## Model

### 1. Description of the system

Our system consists of a population of broiler poultry farmers who purchase, grow, and sell flocks of domestic broiler poultry for the purpose of income. Farmers purchase day-old chicks and sell them as finished birds after a given growing period. Flocks of broiler poultry are kept in coops, and coops may be empty if the opportunity cost of poultry farming is too high. Poultry flocks occupying coops are initially disease-free and can be infected in the course of the growing period if the virus is introduced in the flock. The propagation of the disease in each coop is not modeled; flocks are simply assumed to be infected or uninfected. Coops lose their infected status at the time of depopulation, when infected poultry flocks are sold to traders and replaced by susceptible flocks. Poultry flocks are always sold but the revenue generated by a flock depends on both its growing period (which determines poultry weight) and its infection status (**Supplementary Information 1A**). In a disease-free environment, farmers apply a constant optimal growing period *σ*_Ø_^−1^, and *σ*_Ø_ is the rate of poultry removal (sale) from farms (**Supplementary Information 1A**). Farmers can vaccinate their poultry flocks at an additional cost (**Supplementary Information 1A**), and vaccinated flocks are considered to be fully protected. Each coop is managed by a farmer with knowledge of the infection status of his poultry flocks, and this farmer decides (*i*) whether the coop is populated with poultry or not, (*ii*) whether the flock is vaccinated, and (*iii*) at what age the birds are sold. In a population of farmers, *p̄* is the proportion of coops populated with poultry, with 1−*p̄* coops left empty, and 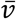 is the proportion of populated coops that are vaccinated.

### 2. Epidemiological model

Coops are divided into categories according to the average sale rate of infected poultry flocks 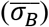. The dynamics of infection among unvaccinated coops in a given category *B* can be written:

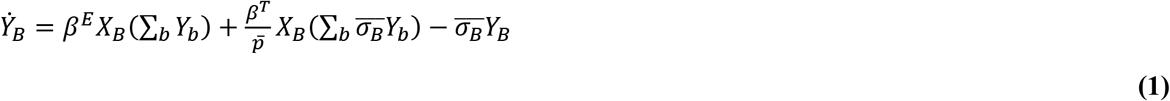

Where *Y*_*B*_ represents the number of infected coops and *X*_*B*_ the number of uninfected coops (see **Supplementary Information 1B** for more detail). The total number of populated coops that are at risk for infection is equal to:

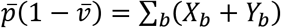

The parameter *β*^*E*^ is the density-dependent environmental transmission term and *β*^*T*^ is the frequency-dependent trade-based transmission term describing farm-to-farm transmission through a network of poultry traders. Environmental transmission (*β*^*E*^) is proximity-based while trade-based transmission (*β*^*T*^) occurs through the trade of infected birds, susceptible coops being contaminated through contact with traders transporting infected poultry (**Supplementary Information 1B**). These two mechanisms are the ones commonly considered as maintaining the circulation of HPAI in countries where the disease is endemic (21). In equation **(1)**, note the 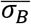 parameter in the second summation corresponding to trade-based transmission; an increased sell rate increases the trade-based force of infection (FOI), as poultry are being moved into the trader network more quickly (**Supplementary Information 1B**).

The basic reproduction number, when *p̄* = 1, 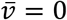 and all farmers apply the same depopulation rate *σ*_Ø_ is 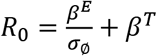

### 3. Economic model

At the individual coop level, a profit optimization takes place. It is assumed that the owner of coop *i* maximizes income generated by the poultry flock in coop *i*. This income is denoted *U*_*i*_

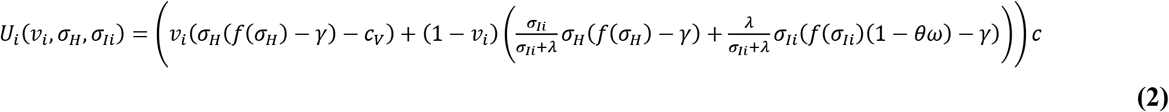

With *v*_*i*_ ∈ [0, 1] the probability that coop *i* is vaccinated, *σ*_*H*_ >0 the sell rate of healthy (either susceptible or vaccinated) flocks, and *σ*_*Ii*_ > 0 the sell rate of infected flocks. The parameters on the right-hand side include *λ* the farm-level force of infection, the parameter *c* ≥ 0 as the unit sale price of poultry meat extracted from healthy birds, and γ the unit replacement cost of flocks (i.e. the purchase of day-old chicks). Without loss of generality, we assume *c* = 1 and *γ* is a fraction of *c*. The parameter *θ* ∈ [0, 1] is the probability that an infected flock is detected as infected at the time of sale, and *ω* ∈ [0, 1] is the proportion decrease in sale price for an infected bird. The product *θω* is hereafter referred to as the *penalty*. We have *c*_*v*_ ∈ [0, 1] as the vaccination cost, defined as a fraction of *c*. *f* is a function relating the sell rate to the carcass weight of slaughtered poultry (**Supplementary Information 1A**). In the remainder of the manuscript, *λ* is considered to be the FOI scaled to *σ*_Ø_^−1^ in other words, *λ*^−1^ represents the number of poultry cohorts cycles that take place before a flock is infected. If *λ*^−1^ < 1, the rate and individual-flock probability of infection are very high. Income flow *U*_*i*_ and all other costs are expressed per growing period *σ*_Ø_^−1^.

A farmer sets his own vaccination status and sell rates (*σ*_*H*_ and *σ*_*I*_). At the population level, stable strategy sets 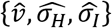 are considered to be game-theoretical stable equilibria if they fulfill the criteria of evolutionary stability (22). Meanwhile the marginal opportunity cost function of poultry farming is considered linear (**Supplementary Information 1C**) and the fraction of coops that are populated is assumed to be:

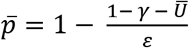

Where 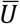 is the average utility across all coops as defined by equation **(2)** and *ε* a constant coefficient. The individual coop parameter *p*_*i*_ ≥ 0 is the probability that coop *i* is populated with poultry, and the linear coefficient *ε* >0 describes the sensitivity of poultry production to changes in income flow.

## Results

### 1. Optimal poultry depopulation rates and epidemiological properties of the system

The optimal value of *σ*_*H*_, by maximizing individual utility in equation **(2)**, can be shown to be independent of *λ* and equal to *σ*_Ø_. Meanwhile *σ*_*I*_ (the sell rate of infected poultry flocks) can be shown to be approximately optimized at the boundary values of 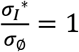 or 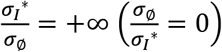 depending on the FOI and penalty (details in in **Supplementary Information 2**). Thus, infected poultry flocks will either be sold when fully grown so that their total revenue can be maximized (this occurs under conditions of a high FOI or low penalty), or they will be sold immediately upon noticing an infection in order to repopulate the coop with a healthy flock (this occurs under conditions of a low FOI or a high penalty). Because of this dichotomous behavior in the sell rate, we allow for two behaviors *D* and Ø, where *D* corresponds to immediate depopulation upon infection and Ø represents the “ null behavior” of waiting until poultry are fully grown before selling the flocks irrespective of infection status (**Figure 1**). Given that a coop is populated and not vaccinated, the individual probability that farmer *i* opts for behavior *D* is *d*_*i*_.

**Figure 1.**
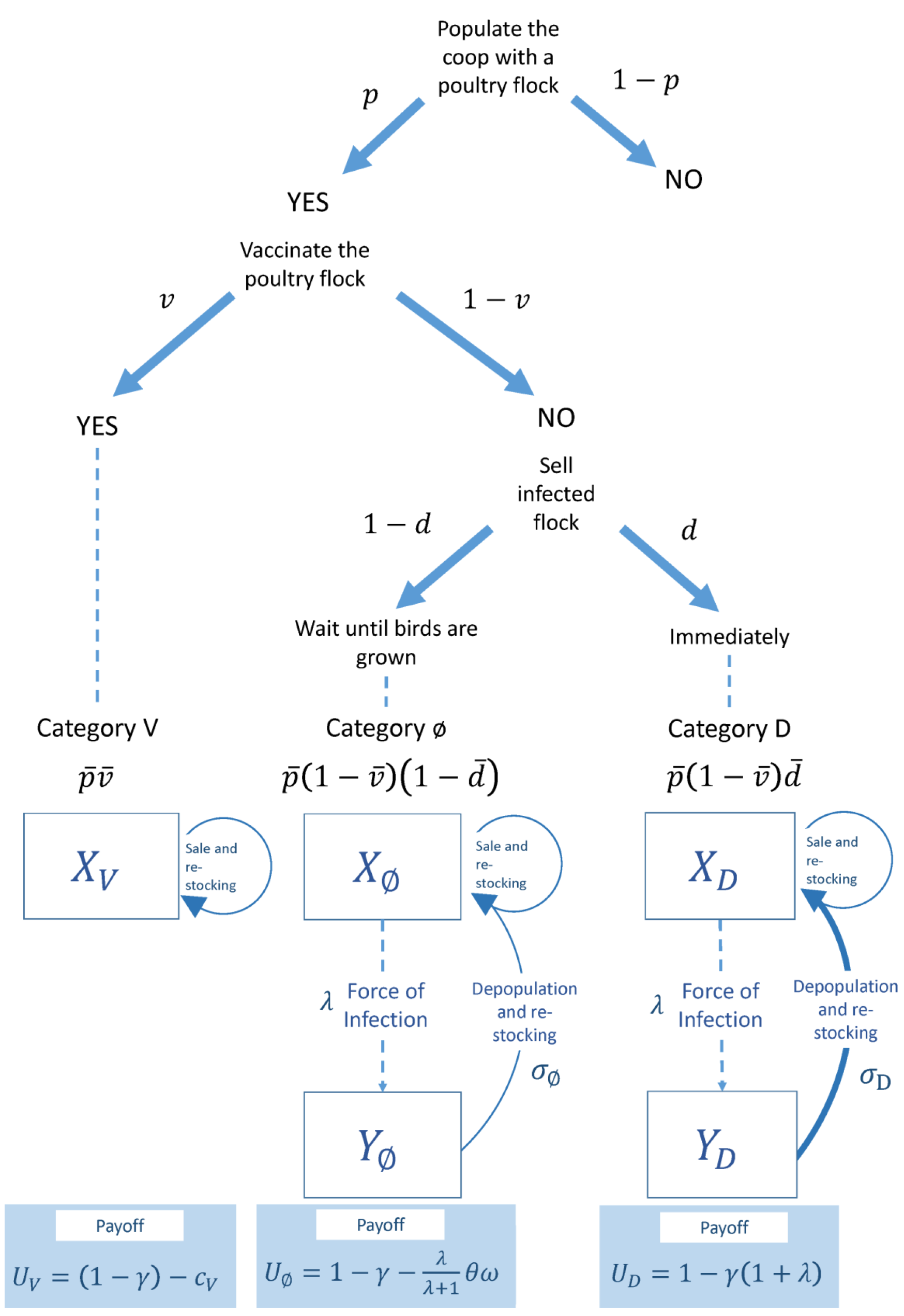
Schematic representation of the farmer’s decision process and associated payoffs for the three management behaviors.

If we assume that the growth period *σ*_*D*_^−1^ in a depopulator strategy is very close to zero 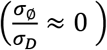 we can define *U*_*V*_ *U*_*D*_ and *U*_Ø_ as the individual payoffs of vaccination, depopulation, and the “null” behavior (see equations in **Figure 1**). Then, the utility function in equation **(2)** simplifies to:

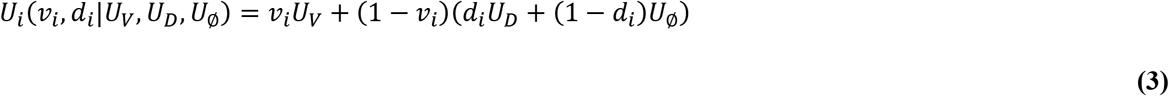

In reality, the sale of infected flocks cannot be simultaneous with infection since it takes some time for farmers to arrange the sale with a trader. However, assuming 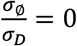 returns values of disease prevalence, *U*_Ø_and *U*_*D*_ which are close to the one obtained with 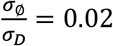 or 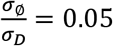, while ensuring mathematical tractability (**Supplementary Information 3**). The ratios 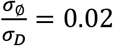 and 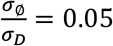 mean that it takes an average of two days or five days, respectively, to sell an infected flock for a standard growing period of 100 days (a common growing period for local and mixed local-exotic breeds of chickens commonly farmed in Southeast Asia) (23).

In general, depopulation and vaccination are incentivized when the penalty is high, as the penalty decreases the payoff for the null behavior Ø. A high FOI tends to favor vaccination (*V*), while low FOI favors depopulation (*D*). From the farmer’s perspective, it is worth the cost to abandon the present revenue from an infected flock and depopulate immediately, if the subsequent flock (initiated immediately after depopulation) is at sufficiently low risk of being infected (**Supplementary Information 2**).

Infected coops in category *D* are not likely to contaminate other coops through environmental transmission because their infectious period is very short. However, since they depopulate almost immediately upon infection, they relay all of their incoming infections into the trader network, and therefore sell a higher absolute number of infected flocks than coops of category Ø. As a result, as 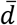 increases, susceptible coops are less likely to be contaminated through environmental transmission from neighboring farms and are more likely to be contaminated through contacts with poultry traders carrying the virus. The dependence of disease prevalence on 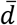 depends on the value of the trade-transmission coefficient *β*^*T*^ (**Figure 2**). If *β*^*T*^ is low enough (*β*^*T*^ < 1), an increase in 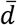 decreases the FOI, eventually resulting in disease eradication; if *β*^*T*^ is high (*β*^*T*^ > 1), an increase in 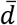 increases the infection risk for both categories of behaviors.

**Figure 2.**
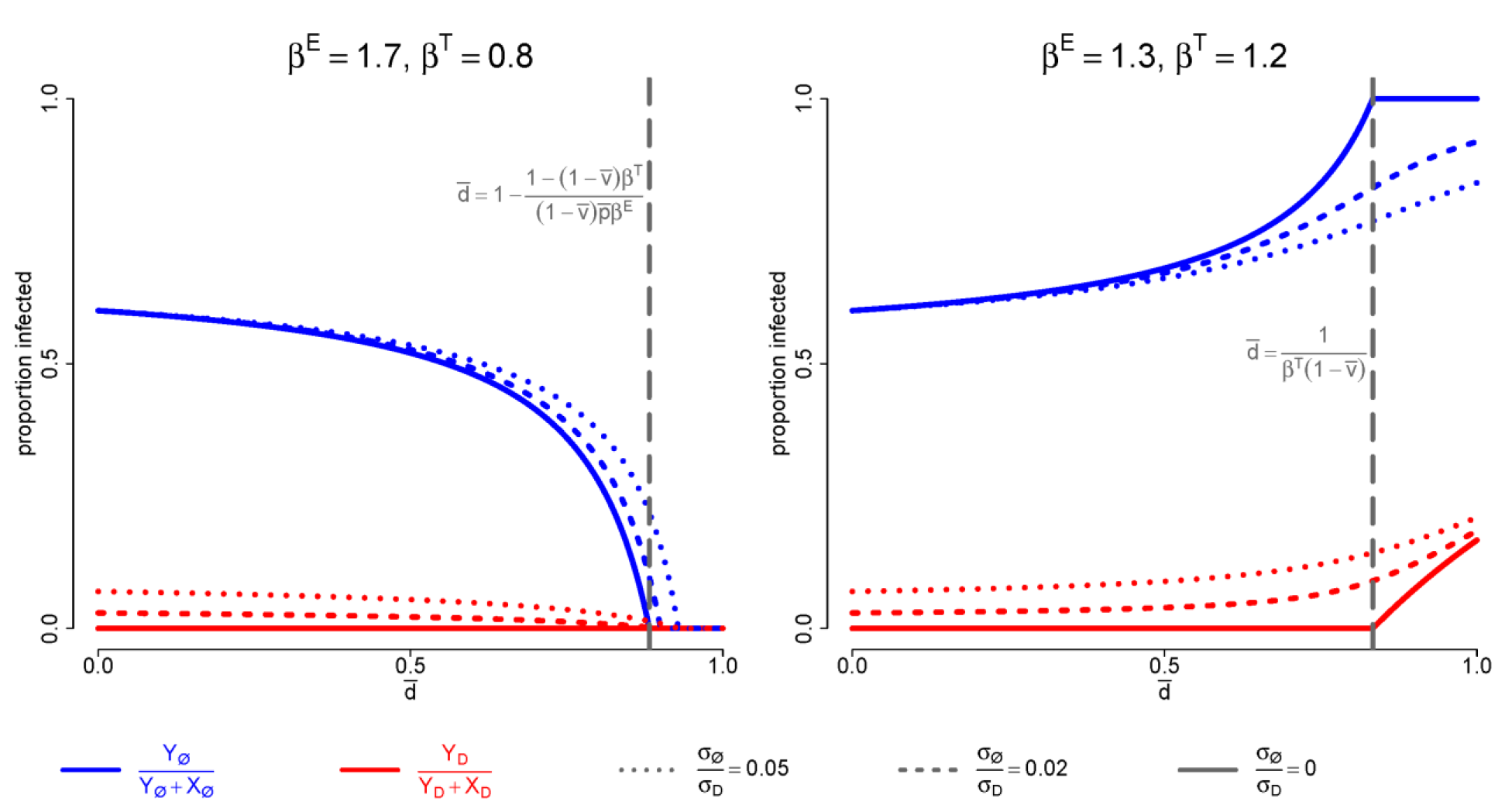
Relationship between the equilibrium proportion of infected coops and proportion of depopulators. In the two panels 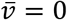, *p̄* = 1, and 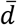 shown on the horizontal axis, with all other variables held constant. The solid lines show the equilibrium prevalence when depopulation is instantaneous, while the dashed and dotted lines show the equilibrium prevalence when depopulated flocks spend 2% (dashed) or 5% (dotted) of their growing period on the farm before depopulation.

### 2. Stability of farmers’ strategies and consequences for disease control

There is a substantial qualitative difference between poultry producer communities where poultry trade alone cannot sustain viral circulation (*β*^*T*^ < 1) (**Figure 3**) and those where it can (*β*^*T*^ > 1) (**Figure 4**). When trade-maintained endemicity is absent (*β*^*T*^ < 1), the incentive created by a penalty on infected poultry allows the depopulation strategy to establish (leading to a stable disease-free equilibrium (DFE)) when either the penalty is sufficiently high or *R*_0_ is sufficiently low. In the presence of trade-maintained endemicity (*β*^*T*^ > 1), the depopulation behavior will persist 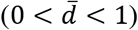 with a low *R*_0_ and intermediate penalty, or fix 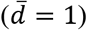 with a high enough penalty, but neither of these scenarios are associated with a stable DFE, as a higher 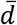 increases the FOI.

**Figure 3.**
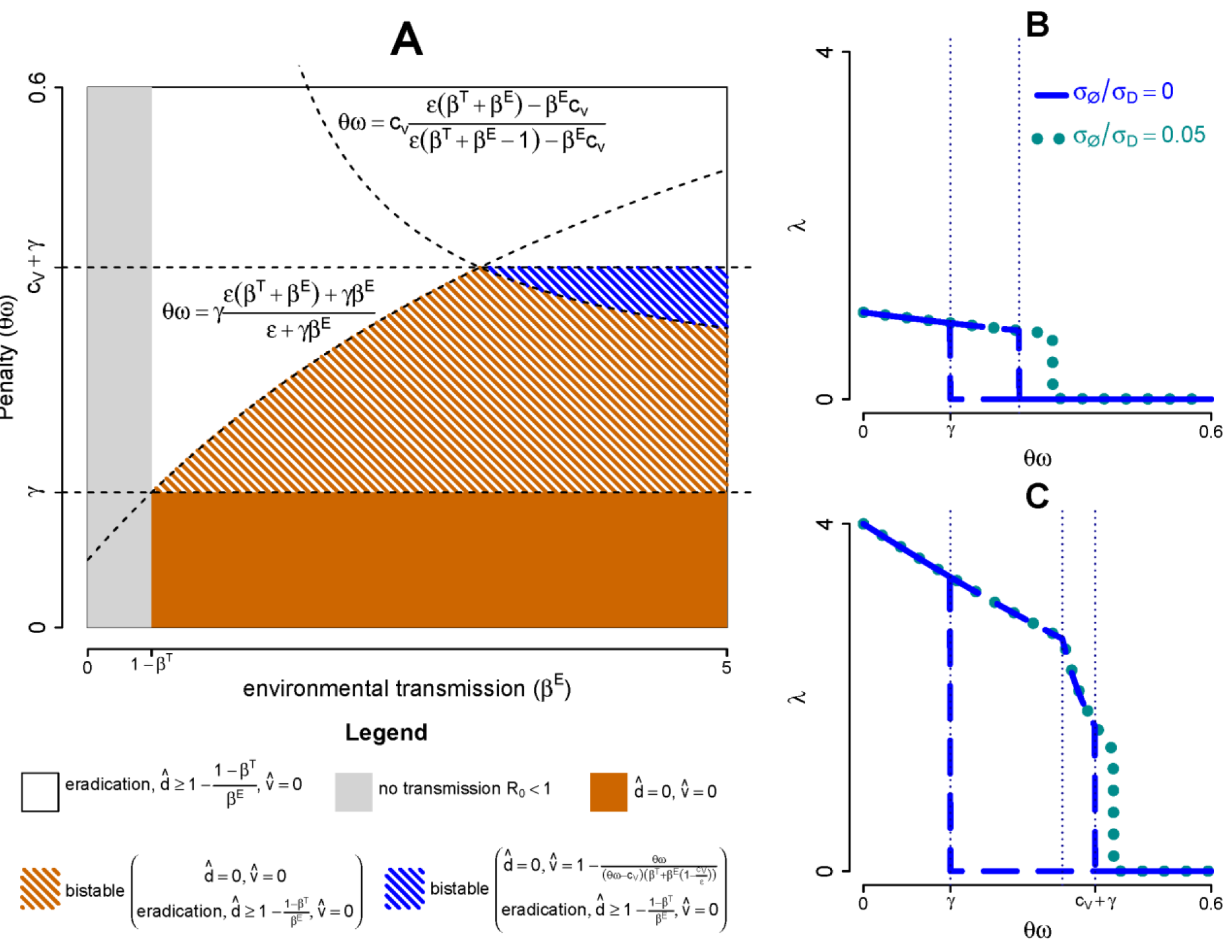
Stable farmer strategies and resulting disease incidence when *β^T^* < 1. (A)predicted stable equilibrium strategies in response to given sets of parameters (environmental transmission, penalty). **(B)** and **(C)**: evolution of the force of infection (*λ*) in response to varying penalty, with *σ*_Ø_/*σ*_*D*_ = 0 and *σ*_Ø_/*σ*_*D*_ = 0.05 (computed numerically) for *R*_0_ = 2 (panel B) and *R*_0_ = 5 (panel C). Parameter values are: *β*^*T*^ = 0.5, γ = 0.15,*c*_*v*_ = 0.25 and *ε* = 0.85. When *R*_0_ is low, increasing the penalty makes the system transit through, successively, an endemic state with pure “null” behavior strategy (Ø), a bistable state of pure Ø or disease-free (with a high proportion of depopulators), and a unique disease-free state. When *R*_0_ is sufficiently high, intermediate values of the penalty result in a mixed strategy of the null behavior and vaccination (*V*, Ø).

In the absence of trade-maintained endemicity, depopulation behavior in the population is self-reinforcing: as 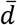 increases, the FOI drops, and the payoff to the depopulation strategy increases as it becomes less likely that a farmer’s subsequent flock will experience infection. This explains the bistability and hysteresis observed in the system with respect to changes in the penalty (**Figure 3**). A high penalty is necessary to eradicate the disease when avian influenza is endemic, but the DFE is maintained when the penalty is subsequently reduced.

In the presence of trade-maintained endemicity, increased adoption of the depopulation behavior is self-defeating: as 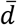 increases, the FOI increases as traders become more likely to contract the infection, and the depopulation strategy’s payoff decreases as the restocking of a coop with a new flock is more likely to lead to reinfection. In this case the mixed strategies (*D*, Ø) and (*D*,*V*) can both be evolutionarily stable, depending on *R*_0_ and the size of the penalty (**Figure 4**). In the mixed (*D*, *V*) strategy, depopulators can be seen as free riders generating perverse epidemiological effects: their depopulation behavior increases in payoff as the number of vaccinators increases, but adopting depopulation behavior (over the null behavior Ø) increases overall disease transmission via the poultry trading network. For the mixed strategy (*D*, Ø), an increased number of depopulators leads to increased FOI, which decreases the payoff of the depopulation strategy (**Figure 4**). In other words, the first depopulator in a ‘pure Ø’ strategy perceives a benefit as the FOI is low, but the marginal payoff decreases as the depopulation behavior is adopted more widely, because more depopulators leads to an increased FOI.

**Figure 4.**
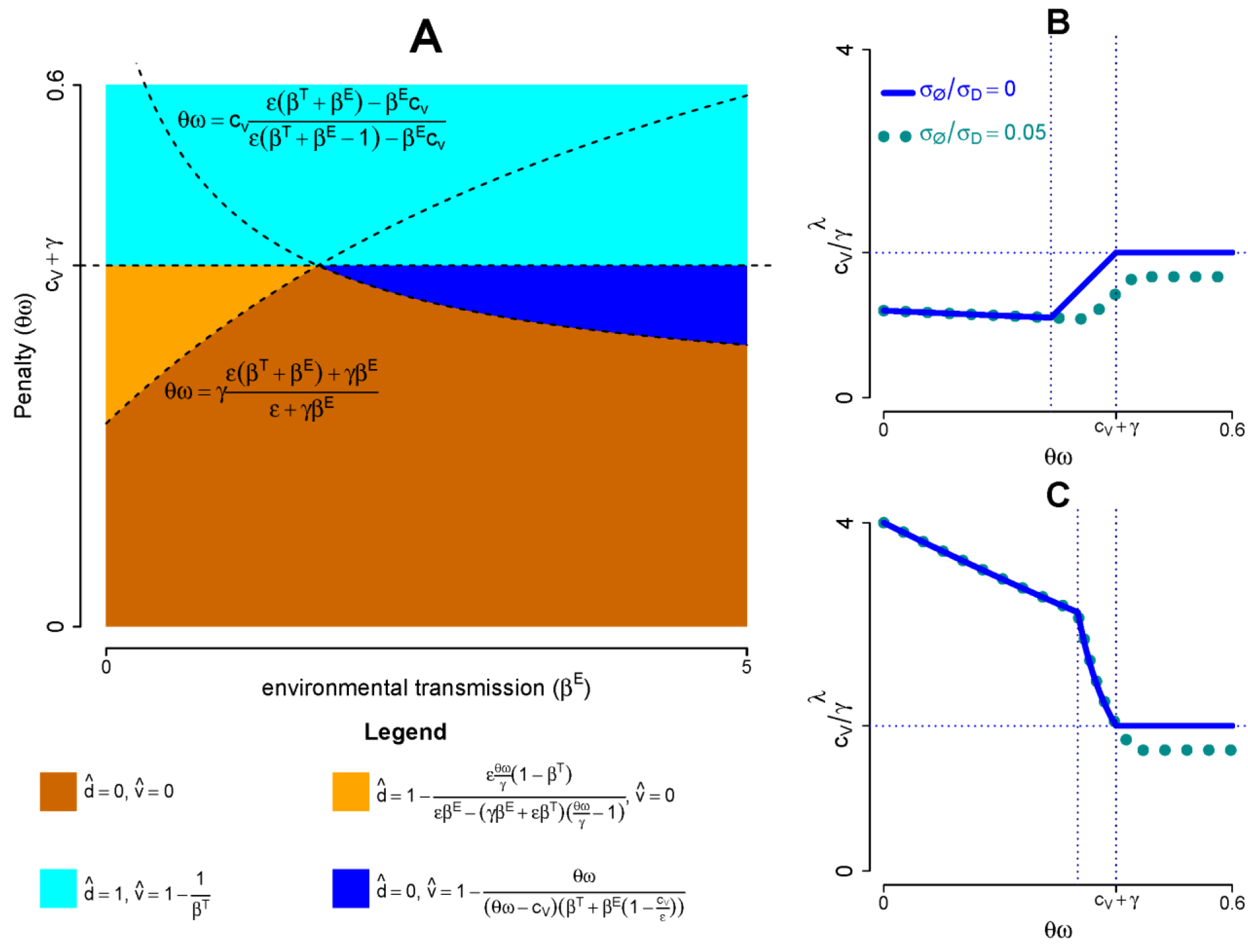
Stable farmer strategies and resulting disease incidence when *β^T^* > 1. **(A)** predicted stable equilibrium strategies in response to given sets of parameters (environmental transmission, penalty). **(B)** and **(C)**: evolution of the force of infection (*λ*) in response to varying penalty, with *σ*_Ø_/*σ*_*D*_ = 0 and *σ*_Ø_/*σ*_*D*_ = 0.05 (computed numerically) for *R*_0_ = 2 (panel B) and *R*_0_ = 5 (panel C). Parameter values are: *β^*T*^* = 0.5, *γ* = 0.15,*c*_*v*_ = 0.25 and *ε* = 0.85. Increasing the penalty on infected flocks makes the system transit from an endemic stable equilibrium where farmers implement the null behavior as a pure strategy to another endemic stable equilibrium where farmers implement either a (*D*, Ø)-strategy for low *R*_0_, or a (*V*, Ø)-strategy for high *R*_0_. If the penalty crosses the threshold *c*_*V*_ + *γ*, farmers will settle into a (*D*, *V*) mixed strategy and the disease will remain endemic.

Vaccination alone cannot maintain a disease-free state since the payoff of vaccination declines as the FOI drops, as demonstrated previously (2, 24). However, the decrease of FOI due to the vaccination of a fraction of the coops incentivizes the rapid depopulation of unvaccinated infected coops, provided the penalty is sufficiently high (*θω* > *c*_*v*_ + *γ*). In the absence of trade-maintained endemicity, this synergistic effect between vaccination and fast depopulation can maintain the disease-free state, even for very high values of *R*_0_, as the presence of this fast depopulating behavior among the non-vaccinees breaks the transmission chain (**Figure 3**). In the case of trade-maintained endemicity, the same synergistic effect maintains an endemic equilibrium - at the game-theoretic evolutionary stable strategy where vaccinators and depopulators co-exist - at which the FOI (equal to 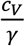) is independent of the individual transmission coefficients and the price penalty (**Figure 4**).

Generally, the penalty only affects the revenue of farmers with the null behavior Ø (**Figure 1**). Thus, as long as 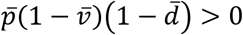 (i.e. all or part of the farmers implement Ø), increasing the penalty disincentives the population of coops with poultry (i.e. decreases *p̄*) and, in turn, decreases the FOI, since the disease transmission is partly density-dependent. However, when all farmers are either vaccinators or depopulators, their revenue is not affected by penalty and the system is irresponsive to changes in penalty (**Figure 4**).

## Discussion

In a multi-player game where a community of farmers seek to maximize their income, we found that the presence of inter-farm poultry trading networks has the largest qualitative effect on the system’s behavior. With low levels of trade-based disease transmission, it is possible to incentivize either depopulation alone or a combination of depopulation and vaccination such that the disease is pushed to an eradication state. With high levels of trade-based disease transmission, adoption of depopulation behavior increases the FOI on poultry traders, and the disease cannot be eradicated. Vaccination does reduce transmission in this scenario, but the lower overall FOI caused by increased vaccination leads to depopulators free-riding on this lower FOI, resulting in an increase of the FOI on traders specifically. As a result, the depopulation behavior cannot be avoided because it is more attractive than vaccination when the FOI is low enough (but not zero). When trade-based transmission alone can sustain endemicity, the disease cannot be eradicated because depopulation persists.

The relevance of the game-theoretical stability to epidemiological-economic systems relies on the assumption of perfect mobility of players: it is assumed that in the long run, actors initially implementing a suboptimal strategy either switch to the optimal one (through adaptation or imitation) or quit the market (i.e. leave their coops unpopulated). This assumption might hold in the context of poultry farming in the developing countries of Southeast Asia, as they are characterized by limited institutional regulation and few barriers to entrance and exit of the sector (25–27). Note that, as the system is subject to the economic and ecological changes affecting HPAI dynamics (fluctuation in market prices and climatic variables) (28), stable equilibriums remain theoretical and should be interpreted as states towards which the system tends to converge.

These results highlight the importance of trade-based disease transmission and modulation of the timing of sale -two real features of smallholder livestock systems – on the epidemiological-economic equilibria of avian influenza circulating in a network of profit-maximizing farmers. Results of a sociological survey in Vietnam suggest that fast depopulation is one of the behavioral responses of farmers to HPAI, as respondents reported an increase in poultry sales during epidemic periods (15). The economic context of poultry farming in some endemic countries is favorable to depopulation in response to disease infection: chick and finished poultry are traded with limited equipment (motorcycle for transportation, storage of poultry at home or in enclosures of live bird markets) (29–31) which limits transaction costs associated with the sale and replacement of flocks. Moreover, the limited sanitary controls and flexibility of the trade networks allow the sale of sick and/or young birds and their use for human consumers or by other livestock farms (python, crocodile, fish) (6, 15, 32–35). The depopulation behavior can explain why avian influenza viruses of the H5 subtype are more likely isolated from poultry sampled in live bird markets (34, 36, 37) than in poultry farms (38–40). A case-control study demonstrated that contacts with broiler poultry traders increase the risk of poultry farm infection with H5N1 HPAI in Vietnam (41) while a time series analysis showed the contribution of time variation of trade activity to the seasonality of HPAI H5N1 (28). Spatial analyses conducted in Indonesia and China also showed that proximity to trade networks is a risk factor of H5N1 HPAI reporting (42, 43). Among factors contributing to infection from trade-based transmission are the high frequency of trader visits to poultry farms and the lack of cleaning and disinfection of traders’ vehicles and equipment. Viral amplification in the trade networks may also occur when poultry from different farms are mixed together at the trader’s house or in live bird markets (44), an effect not accounted for in the present study. To the best of our knowledge, variation in the timing of sale of infected livestock has not been addressed from a theoretical epidemiological or economic perspective. A previous theoretical model of smallholder poultry farm management in response to avian influenza included sell rate as one of the parameters optimized by farmers but without allowing a differential policy for susceptible and infected poultry, which differs significantly from the present study (16).

Thus the study shows the need to elucidate the respective contribution of trade-based and environmental transmission in the circulation of HPAI to design disease control policies. In the absence of trade-maintained endemicity, the hysteretic property of the system implies that there is an opportunity for social planners (i.e. the state, a livestock farming organization, or an integrating private actor) to significantly improve disease control and, in turn, poultry farmers’ welfare. Indeed, temporary costly measures to decrease the FOI (through subsidized mandatory vaccination or poultry mass culling) or to increase the penalty (through sanitary inspections and disease surveillance) may incentivize fast depopulation of infected coops and establish a DFE which is sustained on the long term, provided a small penalty is maintained. Note, however, that mass culling policies applied in outbreak areas, when they are accompanied with financial indemnities, can have the perverse effect of increasing the value of infected flocks and, in turn, disincentivizing depopulation of infected farms and increasing the number of coops with poultry (16).

Eradication is not possible when endemicity is maintained through trade. In this case, increasing the penalty, in the absence of affordable vaccine technology, risks simultaneously increasing the FOI and lowering farmer income, leading to lower poultry production and consumption. Here, two options seem reasonable. One is enhancing disease control and/or biosecurity practices in the network of traders. A second is providing farmers with an affordable vaccine technology in order to maintain immunity in poultry populations and decrease the overall FOI. Policymakers may encourage the creation of trustworthy and sustainable certification schemes ensuring that vaccinated birds are sold at higher prices on the open market (45).

It was assumed here that farmers aim at maximizing an income flow function in coops populated with poultry. A recent field study suggested that poultry farmers’ decision making may be affected by altruistic considerations, risk aversion, time preference, and the influence of other actors in the poultry value chain (15). Farmers are concerned with the welfare of neighboring poultry farmers with whom they have social/family connections. For this reason, they may be more inclined to depopulation than our model predicts, as depopulation would be perceived as reducing local disease transmission. On the other hand, risk aversion may favor vaccination over depopulation. It was shown in Vietnam that poultry farmers cooperate mostly with local feed and chick suppliers to manage poultry diseases, partly because these actors sell feed on credit to farmers, giving them economic influence over their customers (46). Those chick suppliers might perceive the depopulation behavior as advantageous for them as it increases the demand for chicks and limits the local spread of the disease, therefore preserving poultry production in their sale area.

The control of avian influenza – on smallholder farms, in markets, and in trading networks – will remain on the global health agenda as long as certain avian influenza subtypes continue exhibiting high mortality in humans. Identifying the origin of these infections and outbreaks is a critical component of their control. Understanding the relationship between the microeconomics of poultry production and microepidemiology of avian influenza transmission will allow us to develop better tools for the control avian influenza outbreaks in smallholder poultry contexts.

## Acknowledgement

The authors are grateful to Dr Timothy Reluga, from the Department of Mathematics of the Pennsylvania State University, for his useful feedback on the present work.

## Funding

The authors received funding form the Pennsylvania State University.

## References

1. Funk S, Salathe M, Jansen VA. Modelling the influence of human behaviour on the spread of infectious diseases: a review. J R Soc Interface. 2010;7(50):1247–56. doi:10.1098/rsif.2010.0142

2. Bauch CT, Earn DJD. Vaccination and the theory of games. P Natl Acad Sci USA. 2004;101(36):13391–4. doi:10.1073/pnas.0403823101

3. Reluga TC. Game theory of social distancing in response to an epidemic. PLoS Comput Biol. 2010;6(5):e1000793. doi:10.1371/journal.pcbi.1000793

4. Malani A, Laxminarayan R. Incentives for Reporting Infectious Disease Outbreaks. J Hum Resour. 2011;46(1):176–202.

5. Bayham J, Kuminoff NV, Gunn Q, Fenichel EP. Measured voluntary avoidance behaviour during the 2009 A/H1N1 epidemic. Proc Biol Sci. 2015;282(1818):20150814. doi:10.1098/rspb.2015.0814

6. Paul M, Baritaux V, Wongnarkpet S, Poolkhet S, Poolkhet C, Thanapongtharm W, Roger F, et al. Practices associated with Highly Pathogenic Avian Influenza spread in traditional poultry marketing chains: Social and economic perspectives. Acta Trop. 2013;126(1):43–53. doi:10.1016/j.actatropica.2013.01.008

7. Garforth CJ, Bailey AP, Tranter RB. Farmers’ attitudes to disease risk management in England: a comparative analysis of sheep and pig farmers. Prev Vet Med. 2013;110(3/4):456–66.

8. FAO. H5N1 HPAI Global overview-November and december 2010. Rome: Food and Agriclture Organization of the United Nations2010.

9. FAO. EMPRES-I, Global Animal Disease Information System. Food and Agriculture Organization of the United Nations; 2014 [05/05/2015]; Available from: http://empres-i.fao.org/eipws3g/.

10. Gilbert M, Xiao X, Pfeiffer DU, Epprecht M, Boles S, Czarnecki C, Roger F, et al. Mapping H5N1 highly pathogenic avian influenza risk in Southeast Asia. P Natl Acad Sci USA. 2008;105(12):4769–74. doi:10.1073/pnas.0710581105

11. WHO. Avian Influenza. World Health Organization; 2014 [10/10/2015]; Available from: http://www.who.int/mediacentre/factsheets/avian_influenza/en/.

12. Imai M, Watanabe T, Hatta M, Das SC, Ozawa M, Shinya K, Roger F, et al. Experimental adaptation of an influenza H5 HA confers respiratory droplet transmission to a reassortant H5 HA/H1N1 virus in ferrets. Nature. 2012;486(7403):420–8. doi:10.1038/nature10831

13. FAO. Approaches to controlling, preventing and eliminating H5N1 Highly Pathogenic Avian Influenza in endemic countries. Rome: Food and Agriculture Organization of the United Nations; 2011.

14. Sims LD. Lessons learned from Asian H5N1 outbreak control. Avian Dis. 2007;51(1 Suppl):174–81.

15. Delabouglise A, Antoine-Moussiaux N, Phan TD, Dao DC, Nguyen TT, Truong BD, et al. The Perceived Value of Passive Animal Health Surveillance: The Case of Highly Pathogenic Avian Influenza in Vietnam. Zoonoses Public Health. 2016;63(2):112–28. doi:10.1111/zph.12212

16. Boni MF, Galvani AP, Wickelgren AL, Malani A. Economic epidemiology of avian influenza on smallholder poultry farms. Theor Pop Biol. 2013;90:135–44. doi:10.1016/j.tpb.2013.10.001

17. Gramig BM, Horan RD. Jointly determined livestock disease dynamics and decentralised economic behaviour. J Agr Resour Ec. 2011;55(3):393–410. doi:10.1111/j.1467-8489.2011.00543.x

18. Gramig BM, Horan RD, Wolf CA. Livestock disease indemnity design when moral hazard is followed by adverse selection. Am J Agr Econ. 2009:1–15. doi:10.1111/j.1467-8276.2009.01256.x

19. Hennessy DA. Behavioral Incentives, Equilibrium Endemic Disease, and Health Management Policy for Farmed Animals. Am J Agr Econ. 2007;89(3):698–711. doi:10.1111/j.1467-8276.2007.01001.x

20. Beach RH, Poulos C, Pattanayak SK. Farm economics of bird flu. Can J Agr Econ. 2007;55(4):471–83. doi:10.1111/j.1467-8489.2011.00543.x

21. Minh PQ, Stevenson MA, Jewell C, French N, Schauer B. Spatio-temporal analyses of highly pathogenic avian influenza H5N1 outbreaks in the Mekong River Delta, Vietnam, 2009. Spat Spatiotemporal Epidemiol. 2011;2:49–57.

22. Thomas B. Evolutionary Stability: States and Strategies. Theor Pop Biol. 1984;26:49–67.

23. Moula N, Luc DD, Dang PK, Farnir F, Ton VD, Dang VB, et al. The Ri chicken breed and livelihoods in North Vietnam: characterisation and prospects. Journal of Agriculture and Rural Development in the Tropcics and Subtropics. 2011;112(1):57–69.

24. Geoffard P-Y, Philipson T. Disease eradication: private versus public vaccination. Am Econ Rev. 1997;87.

25. ACI. Poultry Sector Rehabilitation Project - Phase I: The Impact of Avian Influenza on Poultry Sector Restructuring and its Socio-economic Effects. Prepared for the Food and Agriculture Organization of the United Nations. Bethesda, Maryland: Agrifood Consulting International; 2006.

26. Burgos S, Hinrichs J, Otte J, Pfeiffer D, Roland-Holst D K. S, et al. Poultry, HPAI and Livelihoods in Cambodia-A Review. 2008. doi:10.1038/nature10831

27. Hong Hanh PT, Burgos S, Roland-Holst D. The Poultry Sector in Viet Nam: Prospects for Smallholder Producers in the Aftermath of the HPAI Crisis. Pro-Poor Livestock Policy Initiative Research Report. Hanoi, Vietnam: Food and Agriculture Organisation of the United Nations; 2007. doi:10.1111/j.1467-8489.2011.00543.x

28. Delabouglise A, Choisy M, Phan TD, Antoine-Moussiaux N, Peyre M, Vu TD, et al. Economic factors influencing zoonotic disease dynamics: demand for poultry meat and seasonal transmission of avian influenza in Vietnam. Scientific Reports. 2017;7(5905). doi:10.1038/s41598-017-06244-6

29. Phan DT, Vu DT, Dogot T, Lebailly P. Financial analysis of poultry commodity chains in Hanoi Suburb, North of Vietnam. In: Hanoi University of Agriculture Francophone Joint University Council (CIUF), editor. Proceedings of Scientific Research Results-Institutional University Cooperation Program 2008-2012. Hanoi, Vietnam: Hanoi University of Agriculture; 2013. p. 101–6.

30. Van Kerkhove MD, Vong S, Guitian J, Holl D, Mangtani P, San S, et al. Poultry movement networks in Cambodia: implications for surveillance and control of highly pathogenic avian influenza (HPAI/H5N1). Vaccine. 2009;27(45):6345–52.doi:10.1016/j.vaccine.2009.05.004

31. Fournie G, Guitian J, Desvaux S, Mangtani P, Ly S, Vu CC, et al. Identifying Live Bird Markets with the Potential to Act as Reservoirs of Avian Influenza A (H5N1) Virus: A Survey in Northern Viet Nam and Cambodia. PLoS One. 2012;7(6):e37986.doi:10.1371/journal.pone.0037986

32. Aust P. An assessment of the commercial production of CITES-listed snake species in Viet Nam and China: IUCN SSC Boa and Python Specialist Group (BPSG)2015.

33. Fearnley L, editor. Disputing Efficacy: Poultry Farmers and Pharmaceutical Exchange in Nanchang County, Jiangxi. 3rd International Workshop on Community-based Data Synthesis, Analysis and Modeling of Highly Pathogenic Avian Influenza H5N1 in Asia; 2011 November 14 – 15 Beijing;China.

34. Phan MQ, Henry W, Bui CB, Do DH, Hoang NV, Thu NT, et al. Detection of HPAI H5N1 viruses in ducks sampled from live bird markets in Vietnam. Epidemiol Infect. 2013;141(3):601–11. doi:10.1017/S0950268812001112

35. Nguyen TTT, Fearnley L, Dinh XT, Tran TTA, Tran TT, Nguyen VT, et al. A Stakeholder Survey on Live Bird Market Closures Policy for Controlling Highly Pathogenic Avian Influenza in Vietnam. Frontiers in Veterinary Science. 2017;4. doi:10.3389/fvets.2017.00136

36. Nguyen DT, Bryant JE, Davis CT, Nguyen LV, Pham LT, Loth L, et al. Prevalence and Distribution of Avian Influenza A(H5N1) Virus Clade Variants in Live Bird Markets of Vietnam, 2011–2013. Avian Diseases. 2014;58(4):599–608. doi:10.1637/10814-030814-Reg

37. Turner JCM, Feeroz MM, Hasan MK, Akhtar S, Walker D, Seiler P, et al. Insight into live bird markets of Bangladesh: an overview of the dynamics of transmission of H5N1 and H9N2 avian influenza viruses. Emerging Microbes & Infections. 2017;6(3):e12–e. doi:10.1038/emi.2016.142

38. Thanh NTL, Vy NHT, Xuyen HTA, Phuong HT, Tuyet PN, Huy NT, et al. No Evidence of On-farm Circulation of Avian Influenza H5 Subtype in Ca Mau Province, Southern Vietnam, March 2016-January 2017. PLoS Current Outbreaks. 2017;9. doi:ecurrents.outbreaks.c816d7333370d68f8a0da33f69168986

39. Desvaux S, Grosbois V, Pham TT, Dao DT, Nguyen TD, Fenwick S, et al. Evaluation of the vaccination efficacy against H5N1 in domestic poultry in the Red River Delta in Vietnam. Epidemiol Infect. 2013;141(4):776–88. doi:10.1017/S0950268812001628

40. Henning J, Henning KA, Morton JM, Long NT, Ha NT, Vu LT, et al. Highly pathogenic avian influenza (H5N1) in ducks and in-contact chickens in backyard and smallholder commercial duck farms in Viet Nam. Prev Vet Med. 2011;101(3–4):229–40. doi:10.1016/j.prevetmed.2010.05.016

41. Desvaux S, Grosbois V, Pham TTH, Fenwick S, Tollis S, Pham NH, et al. Risk Factors of Highly Pathogenic Avian Influenza H5N1 Occurrence at the Village and Farm Levels in the Red River Delta Region in Vietnam. Transbound Emerg Dis. 2011;58(6):492–502. doi:10.1111/j.1865-1682.2011.01227.x

42. Loth L, Gilbert M, Wu J, Czarnecki C, Hidayat M, Xiao X. Identifying risk factors of highly pathogenic avian influenza (H5N1 subtype) in Indonesia. Prev Vet Med. 2011;102(1):50–8. doi:10.1016/j.prevetmed.2011.06.006

43. Fang LQ, de Vlas SJ, Liang S, Looman CW, Gong P, et al. Environmental factors contributing to the spread of H5N1 avian influenza in mainland China. PLoS One. 2008;3(5):e2268. doi:10.1371/journal.pone.0002268

44. Fournié G, Tripodi A, Nguyen TTT, Nguyen VT, Tran TT, Bisson A, et al. Investigating poultry trade patterns to guide avian influenza surveillance and control: a case study in Vietnam. Scientific Reports. 2016;6:29463. doi:10.1038/srep29463

45. Ifft J, Roland-Holst D, Zilberman D. Consumer valuation of safety-labeled free-range chicken: results of a field experiment in Hanoi. Agr Econ. 2012;43(6):607–20. doi:10.1111/j.1574-0862.2012.00607.x

46. Delabouglise A, Dao TH, Truong DB, Nguyen TT, Nguyen NT, Duboz R, et al. When private actors matter: Information-sharing network and surveillance of Highly Pathogenic Avian Influenza in Vietnam. Acta Trop. 2015;147:38–44. doi:10.1016/j.actatropica.2015.03.025

